# Towards Interpretable Cryo-EM: Disentangling Latent Spaces of Molecular Conformations

**DOI:** 10.1101/2024.03.18.585544

**Authors:** David A. Klindt, Aapo Hyvärinen, Axel Levy, Nina Miolane, Frédéric Poitevin

**Affiliations:** LCLS, SLAC National Accelerator Laboratory, Stanford University, CA, USA; Department of Electrical and Computer Engineering, UCSB, CA, USA; Department of Computer Science, University of Helsinki, Finland; Department of Electrical Engineering, Stanford, CA, USA

**Keywords:** cryo-EM, machine learning, ICA, AI for science, disentanglement, physics-based models

## Abstract

Molecules are essential building blocks of life and their different conformations (i.e., shapes) crucially determine the functional role that they play in living organisms. Cryogenic Electron Microscopy (cryo-EM) allows for acquisition of large image datasets of individual molecules. Recent advances in computational cryo-EM have made it possible to learn latent variable models of conformation landscapes. However, interpreting these latent spaces remains a challenge as their individual dimensions are often arbitrary. The key message of our work is that this interpretation challenge can be viewed as an Independent Component Analysis (ICA) problem where we seek models that have the property of identifiability. That means, they have an essentially unique solution, representing a conformational latent space that separates the different degrees of freedom a molecule is equipped with in nature. Thus, we aim to advance the computational field of cryo-EM beyond visualizations as we connect it with the theoretical framework of (nonlinear) ICA and discuss the need for identifiable models, improved metrics, and benchmarks. Moving forward, we propose future directions for enhancing the disentanglement of latent spaces in cryo-EM, refining evaluation metrics and exploring techniques that leverage physics-based decoders of biomolecular systems. Moreover, we discuss how future technological developments in time-resolved single particle imaging may enable the application of nonlinear ICA models that can discover the true conformation changes of molecules in nature. The pursuit of interpretable conformational latent spaces will empower researchers to unravel complex biological processes and facilitate targeted interventions. This has significant implications for drug discovery and structural biology more broadly. More generally, latent variable models are deployed widely across many scientific disciplines. Thus, the argument we present in this work has much broader applications in AI for science if we want to move from impressive nonlinear neural network models to mathematically grounded methods that can help us learn something new about nature.

## 1 INTRODUCTION

Molecules such as proteins or nucleic acids make up the building blocks of life. Living organisms contain a plethora of molecules that often comprise thousands of atoms. Biomolecules change their *conformation* (i.e., shape) to fulfill important biological functions such as enzymatic reactions or cellular communication. Understanding the conformational heterogeneity of biomolecules is crucial for deciphering their functional mechanisms and designing targeted interventions. Cryo-Electron Microscopy (cryo-EM) has emerged as a powerful technique for visualizing molecular structures at high resolution. Recent advancements in computational cryo-EM have demonstrated the potential of latent variable models to capture the diverse conformations adopted by biomolecules (reviewed in Donnat et al., 2022). However, interpreting these learned latent spaces and extracting biologically meaningful information from them remains a significant challenge.

In this paper, we propose a fruitful approach to unravel the complexities of conformational latent spaces in cryo-EM by framing this as an Independent Component Analysis (ICA) problem. A few prior works have tested the application of *linear* ICA to molecular imaging (Gao et al., 2020; Borek et al., 2018) finding more meaningful separation of molecular conformation changes. However, we need theory and models that work for the *nonlinear* models used in modern cryo-EM (e.g. Zhong et al., 2021a). Building on recent theoretical work in identifiable nonlinear ICA Hyvärinen et al. (2024), in disentanglement models and their benchmarks in machine learning Locatello et al. (2019), we suggest a path to bridging the gap between theoretical advancements and practical applications in cryo-EM research. We argue that nonlinear ICA methods have the potential to provide a powerful framework to disentangling the latent representations of biomolecular conformations from cryo-EM datasets, overcoming the limitations of traditional volume visualization approaches and ultimately allowing to delve deeper into the structural dynamics of biomolecules. Moreover, we argue that the establishment of benchmarks and metrics specific to cryo-EM disentanglement models is of paramount importance. Adapting and extending existing benchmarks from the machine learning field should allow to objectively evaluate the performance of different disentanglement approaches and track progress in the development of interpretable cryo-EM methods. Ultimately, this interdisciplinary approach will enhance our understanding of complex biological processes and open up new avenues for therapeutic interventions and drug discovery.

The paper is structured as follows. We first provide a general background on the cryo-EM computational problem (Sec. 2) and how it can be framed as an ICA problem (Sec. 2.1). We then discuss the two fundamental challenges associated with cryo-EM: firstly, separating the conformation and the pose representations (Sec. 2.2) and, secondly, finding the right (disentangled) representation of conformations (Sec. 2.3). We then go into more detail on both challenges by providing quantitative metrics to measure progress and modeling suggestions to improve current frameworks. To disentangle poses and shapes, we propose intervention based metrics (Sec. 3.1) and training schemes (Sec. 3.2) that can be added to existing models. For the larger problem of disentangling conformation representations (Sec. 4.1), we discuss existing disentanglement benchmarks and metrics (Locatello et al., 2019). We then discuss three potential approaches for solving this problem (Sec. 4.2), based on temporal information (Sec. 4.2.1), temperature control (Sec. 4.2.2) and atomic models (Sec. 4.2.3). Finally, we discuss the path forward and the broader implications for this framework to take computational approaches from neural network based curve fitting to actual understanding of the mechanisms in nature.

## 2 INTERPRETABLE HETEROGENEOUS RECONSTRUCTION – A DISENTANGLEMENT PROBLEM

**Table 1.** Glossary. Whenever a distinction is necessary in a given context, we use a * (e.g., *g*^*^) to highlight that we are referring to the *ground truth* model (*g*^*^) or *ground truth* latent variables (*z*^*^, *ϕ*^*^). For instance, we have the *ground truth* generator *g*^*^ of the data, in contrast to the *learned* generator *g* (i.e., decoder) from our model of the data.

| Notation | Description |
| --- | --- |
| $z \in \mathcal{Z} = \mathbb{R}^K$ | Conformation latent variables (vector) |
| $\phi \in \Phi = SO(3)$ | Pose latent variables |
| $g : \mathcal{Z} \times \Phi \rightarrow \mathcal{X}$ | Mixing function / generator / decoder |
| $v \in \mathcal{V} : \mathbb{R}^3 \times \mathcal{Z} \rightarrow \{0, 1\}$ | Volume function (implicit representation) |
| $\pi : \mathcal{V} \times \Phi \rightarrow \mathbb{R}^N$ | Projection / pose function |
| $x \in \mathcal{X} = \mathbb{R}^N$ | Observations (e.g., images) |
| $f : \mathcal{X} \rightarrow \mathcal{Z} \times \Phi$ | Learned representation / encoder |
| ICA | Independent Component Analysis |
| *identifiable* | A model with a unique (set) of solutions |
| *disentangle* | Separate different factors (e.g., pose and shape) |

Heterogeneous cryo-EM reconstruction methods aim to model the different conformations that a molecule may assume (Donnat et al., 2022). For instance, we can think of a molecule with a fixed central structure and two adjustable “arms” (see Fig. 1). Clearly, any conformation that this molecule may assume can be described by providing the position of both arms. Thus, these independently moving parts may be thought of as the fundamental *degrees of freedom* of this molecule’s conformations.

**Figure 1.**
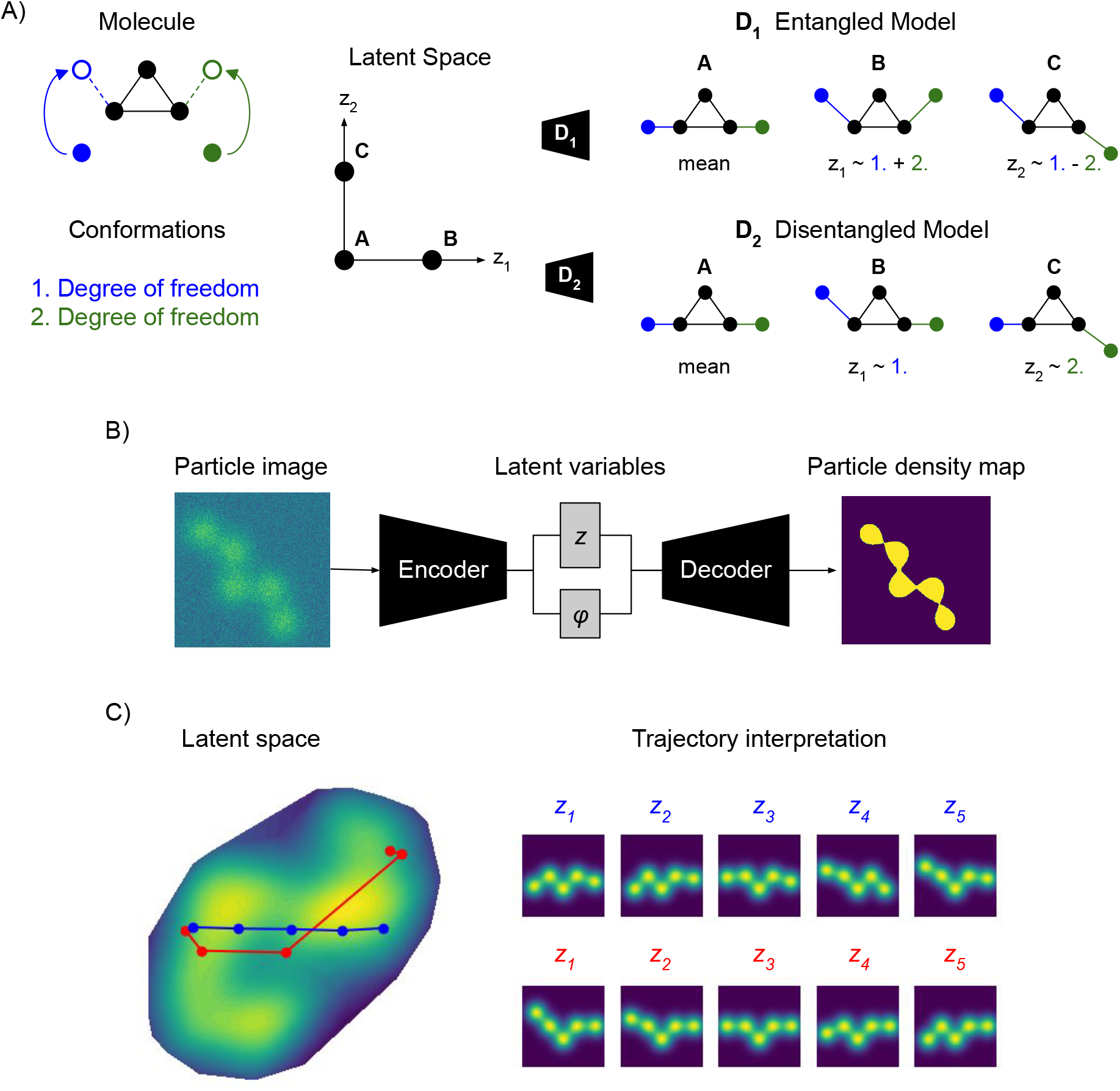
Overview. *What does it mean to have a disentangled representation of molecular conformations?* **A)** The example (left) shows a simple molecule with two degrees of freedom 1. and 2. for changing its conformation. An entangled model (right, top) represents mixtures of both movements on each of its latent dimensions *z*_1_ ∼ 1. + 2. and *z*_2_ ∼ 1. − 2.. A disentangled model (right, bottom) represents pure movements on each of its latent dimensions *z*_1_ ∼ 1. and *z*_2_ ∼ 2.; actually, *z*_2_ = −2. but the sign flip, incorporated in ∼, does not compromise interpretability. Note that disentangling conformations from cryo-EM measurements requires additional information (e.g., time, temperature or physics), as discussed in section 4. **B)** Training a VAE with separate pose *ϕ* and conformation *z* latent spaces on cryo-EM particle images, without any intervention. **C)** Interpreting the learned latent space of a model. An axis traversal (blue) results in a complex motion of both arms, i.e., fails at disentangling the two degrees of freedom. A simple transformation, moving only the left arm, corresponds to a curved trajectory.

We can parameterize them by a two dimensional *latent variable z* ∈ *Ƶ* where *Ƶ* := ℝ^2^ is some degree of movement. The volume that the molecule occupies in three dimensional space can be thought of as a function *v* ∈ *𝒱, v* : ℝ^3^ *×Ƶ* → {0, 1} that is parameterized by *z* and indicates for any position in space (R^3^) whether it is part of the molecular volume or not, known as an *implicit representation* of the volume (Sitzmann et al., 2020; Donnat et al., 2022). That means, for different values of *z, v*_*z*_ = *v*(., *z*) would describe a different volume. Crucially, this function is not known and it is a central goal in heterogeneous cryo-EM reconstruction to learn and study it. For instance, one approach would be to train a neural network (*v*_*θ*_) to approximate the true volume function *v*_*θ*_ ≈ *v*^*^ (Zhong et al., 2021a).

Furthermore, in cryo-EM we typically see a projection *π* : *𝒱 ×* Φ → ℝ^*N*^ to a gray-scale pixel image (represented, to keep notation uncluttered, as a vector with *N* entries). This projection depends on the pose parameters *ϕ* ∈ Φ = *SO*(3), so we will also refer to *π* as the *pose function* (Glossary 1). The pose parameters may also need to be inferred (typically, the cryo-EM image formation model would also include camera parameters such as the microscope defocus — we are skipping those for simplicity). That means, for different values of *ϕ, π*_*ϕ*_ = *π*(., *ϕ*) would describe a different projection. Importantly, the function *π* does not have to be learned because we know the physics, i.e., optics behind this projection, thus, we know that the projection in our model *π*_*ϕ*_ must be the same one as the *ground truth* projection 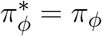.

Putting this together we can write the combined cryo-EM generative model (i.e., the abstract process that yields the data we observe). That is, the observed data *x* is modeled as being generated by the *ground truth* model as *x* = *g*^*^(*z*^*^, *ϕ*^*^), which is, crucially, a function of the *ground truth* latent quantities (*z*^*^, *ϕ*^*^)

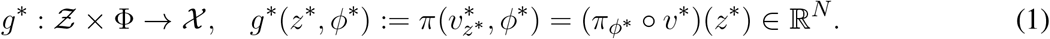

Usually, we would be measuring very noisy signals *x* = *g*^*^(*z*^*^, *ϕ*^*^) + *ϵ* where the noise can be modeled as additive Gaussian *ϵ* ∼ *𝒩* (0, *σ*^2^) in image space. The essential problem of heterogeneous reconstruction in computational cryo-EM can now be stated as follows.

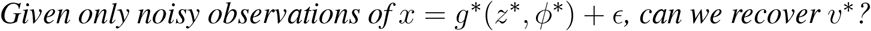

This would be the true volume function *v*^*^ that shows us how the independent degrees of freedom change the molecule’s conformation. Many cryo-EM models actually learn a probabilistic *p*(*x*|*z, ϕ*) representation of the observed data *x* conditioned on the latent variables. Thus, it becomes necessary to perform inference such as maximum *a posteriori* estimation of the latents, conditioned on some observed data *p*(*z, ϕ*|*x*). Alternatively, it is common to approximate the posterior distribution itself with an amortized variational method such as a variational autoencoder (VAE) (Kingma and Welling, 2013). In this work we will be agnostic about the inference procedure (maximum *a posteriori* probability (MAP), or mean of the amortized variational posterior) and just assume that there exists a mapping *f* : *𝒳* → *Ƶ* from data to latent variables.

### 2.1 Heterogeneous Reconstruction in Cryo-EM is an ICA Problem

Let us compare this to a standard independent component analysis (ICA) setting (Comon, 1994). In ICA we assume that there are *K >* 1 independent variables collected in the random vector *z* = (*z*_1_, …, *z*_*K*_). As an example to illustrate this, we may think of a public space where *K* different speakers proclaim their prophecies *z*_*i*_, completely independently of one another *p*(*z*_*i*_, *z*_*j*_) = *p*(*z*_*i*_)*p*(*z*_*j*_), ∀*i*≠ *j*. However, we do not observe those *z* directly. Instead, we observe *K* linear combinations of those variables

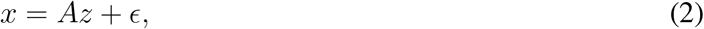

with *A* ∈ *ℛ*^*K×K*^ some *unknown* full rank matrix and, again, with additive Gaussian *ϵ* ∼ *𝒩* (0, *σ*^2^). In our example, this may correspond to *K* microphones placed in the space and each recording some linear combination 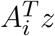 of the speech signals. This scenario is also called *blind source separation*, the term “blind” referring to the idea that we know almost nothing about the “sources” *z*_*i*_, apart from some general statistical properties. In linear ICA, the function *g* : *Ƶ* → *𝒳, g*(*z*) = *Az* that maps from *sources z* to *observations x* is called the *mixing function*. This basic case of linear ICA has been well-studied in the machine learning and signal processing literature (Hyvärinen and Oja, 2000). Briefly, under the simple assumption that at most one of the sources *z*_*i*_ follows a Gaussian distribution, we can find an *unmixing function f* : *𝒳* → *Ƶ* that approximately inverts the mixing function, in practice up to (*f* ◦ *g*)(*z*) ∼_*C*_ *z*, i.e., some simple equivalence class ∼_*C*_ such as permutations and scalings.^1^

If *g*^*^(*z*^*^, *ϕ*) in eq. 1 was linear in *z* and *ϕ*, then the cryo-EM reconstruction problem would amount to a simple linear ICA problem with the (extended) sources (*z*||*ϕ*) where || denotes concatenation. Unfortunately, the cryo-EM mixing function *g*(*z, ϕ*) in eq. 1 is nonlinear. This can be easily appreciated, e.g., by noting that a multiple of some latent *αz* will not produce the same output as an equally scaled image *g*(*αz, ϕ*) ≠ *αg*(*z, ϕ*) which would be just the same image but changed in brightness. In the case of a nonlinear mixing function *g*, Hyvärinen and Pajunen (1999) showed that it is possible to construct many functions *f* : *𝒳* → *Ƶ* that turn the data into independent variables. However, most of these independent variables have no intelligible relationship with the true sources *z*. This problem is called the lack of *identifiability* of the model, which in general mathematical terms means lack of uniqueness of the solution.

For cryo-EM models this would mean that we can learn *latent spaces* whose individual dimensions have no principled relationship with the true degrees of freedom in molecular conformations. As an example, we may end up with a representation of the simple two dimensional molecule from above where traversing any single dimension in the latent space of our model corresponds to complex combinations of the two arm movements (Fig. 1). This would, likely, bias our interpretation of how they are articulated together to carry out their function. Thus, without further restrictions on our model, we would fail to discover the simple and elegant structure where the molecule just changes conformation along two independent degrees of freedom, i.e., left and right arm. In the modern machine learning context, finding a latent space that separates the underlying factors of variation is often called *disentanglement* (Bengio et al., 2013), but it has to be noted that the meaning of that term is quite vague.

Fortunately, recent advances propose ways to solve this problem with nonlinear ICA (Hyvärinen et al., 2024). For example, Khemakhem et al. (2020) adds conditioning (“auxiliary”) variables *u* that change the source distributions *p*(*z*_*i*_|*u*). Such a *u* could represent extra measurements by another modality, or it could be defined by interventions. The model then becomes identifiable if the *u* modulates the distribution of *z* strongly enough. This is possible because then the *z*_*i*_ are conditionally independent for any *u*, which provides much stronger constraints than the mere (unconditional) independence of the *z*_*i*_ as in the basic ICA framework. Khemakhem et al. (2020) further propose to estimate this model using variational methods, leading to an algorithm which is a variant of VAEs. An alternative approach is possible by assuming temporal dependencies of the source time series (Hyvarinen and Morioka, 2017; Klindt et al., 2020; Hälvä et al., 2021); spatial dependencies can also be used (Hälvä et al., 2024). In this case, independence of the components over time lags leads, again, to more constraints, and thus to identifiability under some conditions. A very different approach can be developed by constraining the nonlinear function *g*, parameterizing it with such a small number of parameters that identifiability is obtained (Hyvärinen et al., 2024, Sec. 5.4); for example, if we know the physics underlying *g* (i.e., pose transformations and projections) we may also be able to obtain identifiability. Finally, we point out that the independence assumption can be relaxed (Träuble et al., 2021); even causal relationships between the independent components have been modeled, but this requires further constraints and assumptions (Träuble et al., 2021; Morioka and Hyvärinen, 2023; Yao et al., 2023). Any such learning is easier if interventions on the system are possible (Ahuja et al., 2023) or if it is assumed that the system undergoes sparse, discrete state changes like in robotics experiments (Locatello et al., 2020b), but the theory mentioned above is specifically *unsupervised*, thus not necessarily requiring interventions.

In this work, we will argue that *the heterogeneous reconstruction problem in cryo-EM should be framed as a non-linear ICA problem* to help us build better and more interpretable models that separate the independent degrees of freedom with which molecules change conformation in nature. Few prior works have applied *linear* ICA to molecular imaging (Gao et al., 2020; Borek et al., 2018), finding more meaningful separation of molecular conformation changes (Fig. 2). However, to the best of our knowledge, none of the nonlinear ICA approaches mentioned above have so far been applied to the field of computational cryo-EM. Below we will propose three promising candidate approaches for solving the nonlinear ICA problem in cryo-EM.

**Figure 2.**
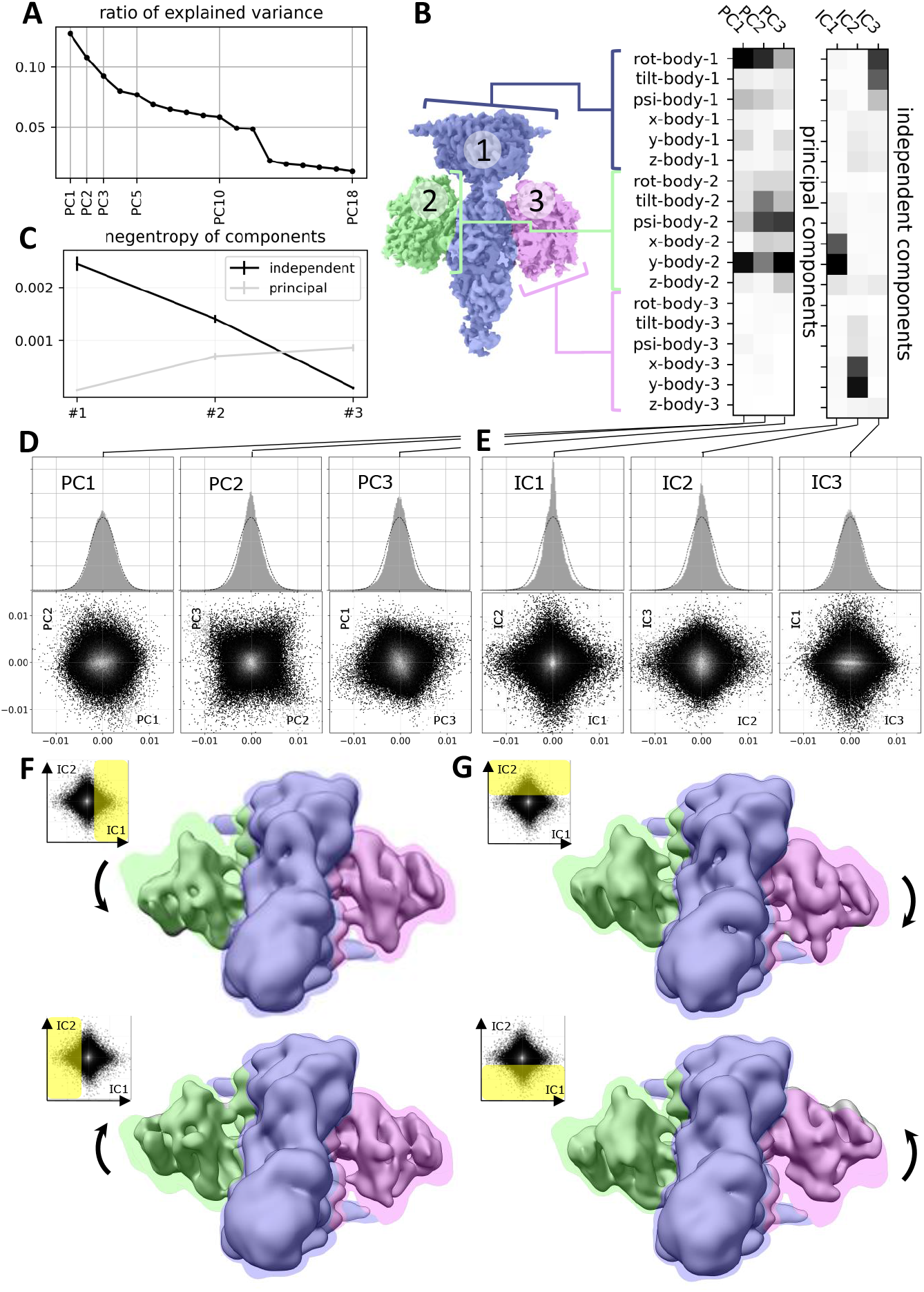
Application of ICA to cryo-EM data. (reproduced with permission from (Gao et al., 2020)). Original caption: **A)** Principal component analysis of the 18 multi-body parameters refined for each particle image yields 18 principal components (PC) displayed here in decreasing order of explained variance. The first 3 components explain more than 30% of the variability in the particle images. **B)** (left) definition of the multi-body segmentation: the central PDE6 stalk in blue corresponds to Body 1, while the two G*α*T·GTP subunits correspond to Bodies 2 and 3. (right) The motion of each body is parameterized with 3 translational parameters and 3 rotational parameters. Each of the 18 principal and 3 independent components is a linear combination of the resulting 18 rigid-body parameters, and their weights are shown here for the first 3 principal components (from negligible to larger weight as the shade of grey becomes darker). **C)** Negentropy (i.e. reverse entropy) of the first 3 principal and independent components. **D)** (resp. **E)**) – (top) histogram of the projection of all image particle parameters on the first 3 principal (resp. independent) components PC1, PC2 and PC3 (resp. IC1, IC2 and IC3). (bottom) 2D histograms of the projections of all image particle parameters on all pairs of the first 3 principal (resp. independent) components. **F)** (resp. **G)** – Maps illustrating the motions carried by IC1 (resp. IC2). (top) map reconstructed from the particles whose projections belong to the last bin along IC1 (resp. IC2). (bottom) map reconstructed from the particles whose projections belong to the first bin along IC1 (resp. IC2). All maps are shown overlaid on the consensus map, with threshold set at a lower density value, colored according to the scheme in B.

### 2.2 Disentangling Pose and Conformation

The first challenge in computational cryo-EM is that of separating the pose *ϕ* of a molecule from its conformation *z*. Again, looking at the cryo-EM mixing function (eq. 1) *x* = *g*^*^(*z*^*^, *ϕ*^*^) + *ϵ*, this means we want to find a representation

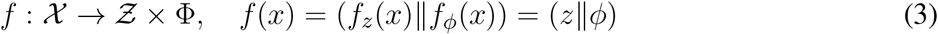

that separates the estimated conformation *z* and the pose *ϕ*.^2^ In other words, as a first step, we could find just two latent *subspaces* without specifying the individual components, or the bases, inside those subspaces. Again, if *g*^*^ were linear, we might use the well-developed methods of independent subspace analysis, or subspace ICA, to approach this problem (Hyvärinen and Hoyer, 2000; Theis, 2006). This might also help with the pose variables’ topology that is, typically, not Euclidean. For instance, a circular pose variable *ϕ*_*i*_ ∈ *S*^1^ that lives on a circle and encodes rotations around one axis could not be represented, by a single dimension, in a typical latent variable model that maps to real valued scalars *f* : *𝒳* → ℝ^*K*^. However, some subspace variants of ICA provide exactly such a transformation into spherical coordinates (Hyvärinen et al., 2009, Ch. 10).

Many cryo-EM models use separate latent spaces to represent conformation and pose (Donnat et al., 2022). However, that does not mean that models learn, during nonlinear optimization, to actually use those separate spaces in the intended way. Recent work by Edelberg and Lederman (2023) demonstrated that this is a problem in popular cryo-EM models such as CryoDRGN (Zhong et al., 2021a). In particular, they showed that a 90^◦^ rotation of an image causes a different prediction in the space of conformation latent variables, even though those should be *invariant* to pose transformations (see below, 3.1).

### 2.3 Disentangling Independent Factors of Conformations

A more fundamental challenge is that of separating the independent degrees of freedom of a molecule. Specifically, we want to find a representation *f*_*z*_ of the molecular conformation that inverts, up to some equivalence class ∼_*C*_ like permutations and scaling (see above), the *ground truth* generative model (*f*_*z*_ ◦ *g*^*^)(*z*) = *z* ∼_*C*_ *z*^*^.

A popular approach (see [CryoDRGN tutorial](https://ez-lab.gitbook.io/cryodrgn/cryodrgn-empiar-10076-tutorial)) consists in fitting a nonlinear model to cryo-EM data (Fig. 1B) followed by manual investigation of the learned latent space that represents conformational heterogeneity (Zhong et al., 2021a) (Fig. 1C), thus limiting our ability to quantitatively compare models. Here, we propose a possible remedy in the shape of benchmarks where we simulate data using the generative model (eq. 1) to assess how close different methods get to the *correct* (i.e., up to ∼_*C*_) representation of conformational latent spaces. This taps into a rich, recent literature in nonlinear ICA methods (Hyvärinen et al., 2024) including benchmarks and metrics for model comparisons (Locatello et al., 2020a).

Once we have benchmarks and metrics, we can measure quantitative progress. However, none of the existing heterogeneous reconstruction approaches in computational cryo-EM are identifiable — mirroring the state of the disentangled representation machine learning field in 2019 (Locatello et al., 2019). To actually make progress, in this perspective, we propose three potential approaches to apply nonlinear ICA method for the unsupervised discovery of molecular conformational changes:

#### 1. Time-resolved single particle imaging

Observing conformational changes over time, such as a sparse change in a single conformational degree of freedom, provides valuable information; this relies on nonlinear ICA methods that use temporal autocorrelations of the sources (Sec. 4.2.1).

#### 2. Boltzmann ICA

It may be possible to disentangle conformational degrees of freedom by sampling at different temperatures; this relies on nonlinear ICA methods that use additional conditioning variables *u*, like temperature (Sec. 4.2.2).

#### 3. Atomic models

Building constraint models, with knowledge of the physical mechanism, may exclude faulty solutions; this relies on reducing the hypothesis class provided by the (typically over-parameterized) neural network models (Sec. 4.2.3).

## 3. DISENTANGLING POSE AND CONFORMATION

In this section we will first discuss the problem of separating pose and conformation in cryo-EM latent variable models. Recent experiments by Edelberg and Lederman (2023) demonstrated that this desired disentanglement is, unfortunately, violated in the case of CryoDRGN (Zhong et al., 2021a). We start by proposing more systematic evaluations and metrics to measure progress on this task (Sec. 3.1). These metrics inspire simple supervised intervention experiments that can be executed in simulation and added to existing training pipelines to disentangle pose and conformation in cryo-EM latent spaces (Sec. 3.2).

### 3.1 Evaluating Disentanglement of Pose and Conformation

We are interested in the cryo-EM *ground truth* generative function 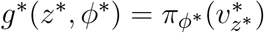 (eq. 1), which consists of a *known* pose function 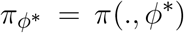, an *unknown* volume function 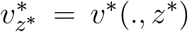, an *unknown* pose *ϕ*^*^ and an *unknown* conformation *z*^*^. Now, for any specific image, we have full control over the pose function 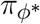 but do not know the pose *ϕ*^*^; however, for the conformation we have neither control over the volume function 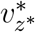 nor knowledge of the *true* conformation *z*^*^ (Shannon et al., 1959). Consequently, in this section and in section 3.2 we leverage the fact that we have complete knowledge about the pose function, to measure and constrain the flexibility of the conformation *z* and volume function *v*_*z*_ that we learn in our model.

Put simply, what we want is that, for a molecule with fixed conformation, our model predicts the same conformation even if we change the pose of the molecule. That is, we want the conformation representation *f*_*z*_ to be *invariant* to pose changes. Additionally, we want the pose representation *f*_*ϕ*_ to be invariant to conformation changes. Mathematically, the requirements of invariance can be written as

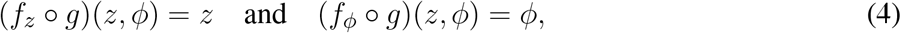

for all possible poses *ϕ* ∈ Φ and conformations *z* ∈ *Z*. When we train a model, this can go wrong both in our *encoder f* (*x*) (if it fails to separate pose and conformation), or in our *decoder g*(*z, ϕ*) = *π*(*v*_*z*_, *ϕ*) (if the volume function *v*_*z*_ learns to represent pose changes). Moreover, it may be necessary to add observation noise to the generated images *g*(*z, ϕ*) + *ϵ* to mitigate for domain shift between the training data and these simulations. To measure progress in this challenge, we can turn this into six different evaluation metrics. We introduce those six in App. 5.1. In practice, a single metric (Alg. 1, eq. 5) seems to suffice as we will discuss in the next section.

### 3.2 Correcting Disentanglement of Pose and Conformation

Based on the ideas proposed in the previous section and the metrics in App. 5.1, we are now going to propose a simple penalty term that can be added to existing cryo-EM models to disentangle pose and conformation. The logic behind these intervention experiments is illustrated in Fig. 3. This procedure relies on the physics-based decoder *g* with an explicit pose representation *π*_*ϕ*_. Typically, the representation 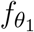

**Figure 3.**
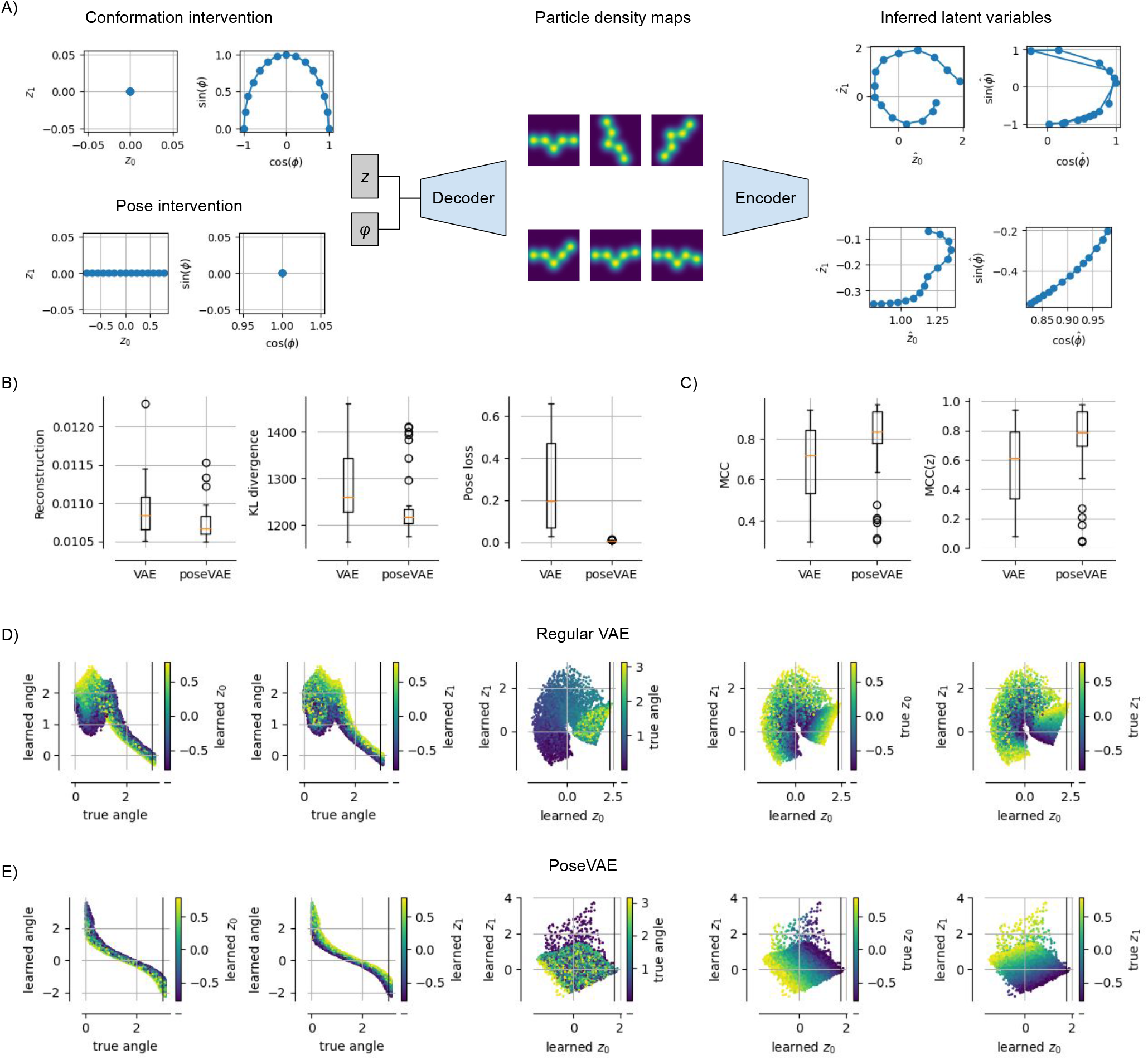
Physics-based Pose and Conformation Disentanglement. **A)** We perform an intervention, i.e., changing only the pose (conformation) of a latent pair; these changed latents are decoded and encoded again to measure the consistency and invariance of our model. **B)** Compared to a vanilla VAE, a model `PoseVAE` trained with interventions (i.e., Alg. 1) achieves lower reconstruction error, lower KL divergence to the prior and a lower pose disentanglement loss (eq. 5). **C)** Disentanglement, measured as mean correlation coefficient (MCC, (Hyvarinen and Morioka, 2017)), increases not only between pose and conformation variables (left), but also among the conformation variables (right). **D)** Visualizations of the learned latents for the vanilla VAE model, showing that the learned angle is not perfectly representing the true angle (plots one and two from the left); the three plots on the right show the learned conformation latents, representing mixtures of true conformation and pose. **E)** Same as **D)** but for `PoseVAE`, showing a perfect monotonic relationship between learned and true angle; also the conformation latents contain little information about the true angle (third plot) and disentangle the true conformation variables up to a 45^◦^ rotation.

#### Algorithm 1: Interventions for Pose and Conformation Disentanglement

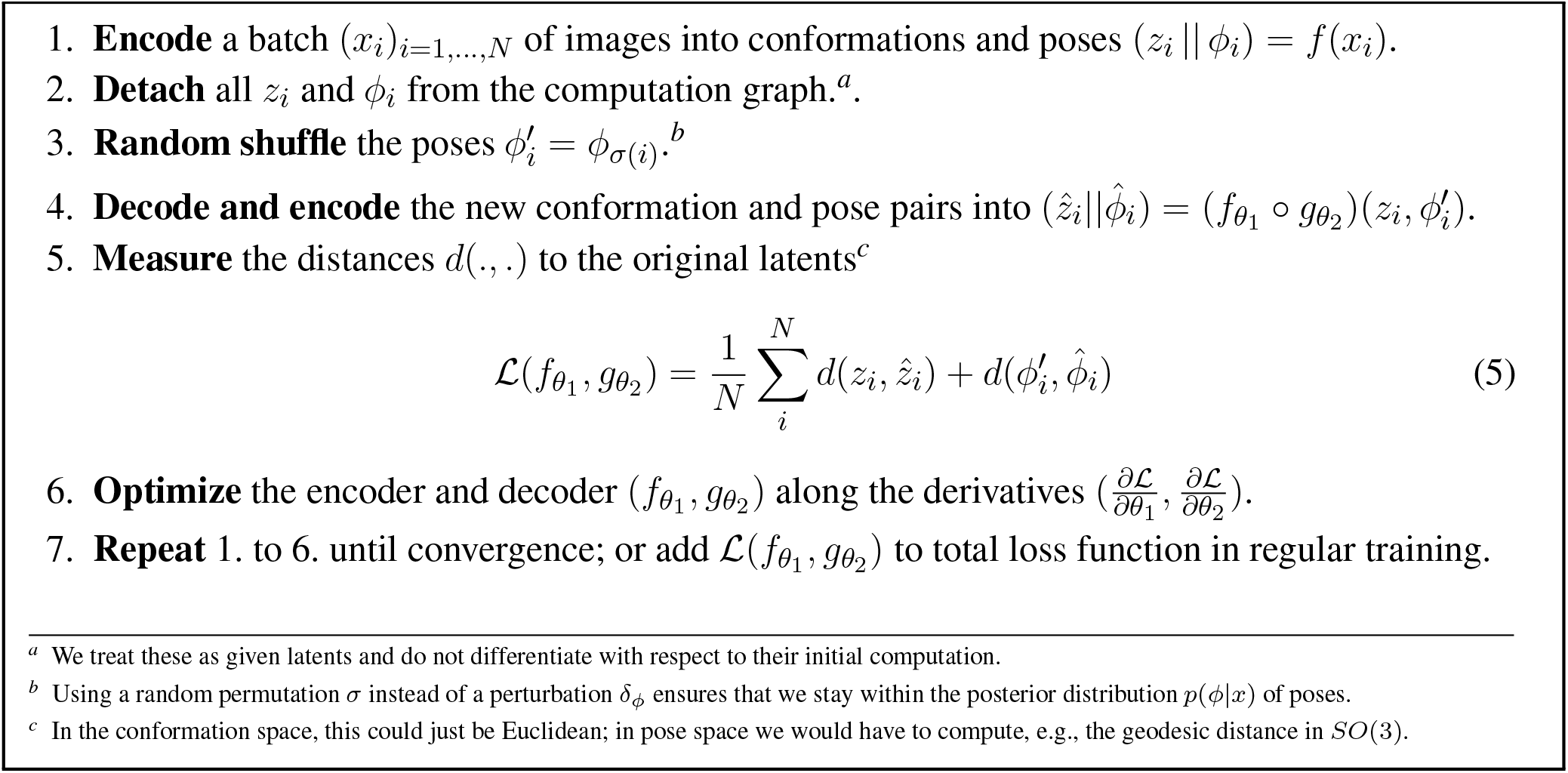

and the generator 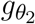 are parameterized as neural networks with learnable parameters *θ*_1_ and *θ*_2_ (Kingma and Welling, 2013; Zhong et al., 2021a). Clearly, we can compute the gradients of all metrics with respect to those parameters. In practice, we observed that good results can be achieved simply by following Alg. 1. Note that this is a straightforward, supervised learning objective that is a relatively standard problem in modern machine learning and should present little difficulty. Thus, we can just add 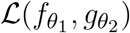 (Alg. 1) as an additional penalty term to the loss function of any of the existing models with separate pose and conformation representation to encourage disentanglement.

Importantly, we are only able to write this approach in such a concise and easy form because of the physics-based decoder *g*. By this we mean the fact that we know the physics, i.e., optics behind the projection *π*_*ϕ*_ in the image formation model (eq. 1). We could imagine a different cryo-EM generative model where both the conformation and the pose change are modeled by the implicit volume representation *v* : ℝ^3^ *× Ƶ*^*′*^ with some extended latent space *Ƶ*^*′*^. Or, in even more general terms, we could just train a standard VAE (Kingma and Welling, 2013) on cryo-EM images to learn a neural network encoder *f* : *𝒳* → *Ƶ*^*′′*^ and decoder *g* : *Ƶ*^*′′*^ → *𝒳* back to image space with some, potentially, even more abstract latent space *Ƶ*^*′′*^ Miolane et al. (2020). However, such abstract models would not have the built-in physics of objects in space, their poses *ϕ* and their projections *π*_*ϕ*_ onto a two dimensional image, which we assume *a priori* in our standard cryo-EM decoder (eq. 1). In other words, such more abstract models would lack the architectural distinction between *z* and *ϕ* which we need in our intervention experiment to disentangle pose and conformation. Thus, we would not be able to manipulate distinct parts of the extended latent spaces (*Ƶ*^*′*^ or *Ƶ*^*′′*^), knowing that those represent distinct physical manipulations of the image.

For models using an implicit representation of the volume, the reason we have to use this interventional approach to disentangle pose and conformation in the first place is that the implicit representation *v*(*z*) is a highly flexible neural network that can easily model pose changes (Sitzmann et al., 2020). Only by combining this with the constraints physics (i.e., the image formation optics) are we able to disentangle pose and conformation representation. This is akin to disentanglement approaches that use the assumption of sparse manipulations, i.e., pairs of data points where only subsets of the latents are modified (Locatello et al., 2020b). Those models have been demonstrated to solve the nonlinear ICA problem theoretically and practically. Thus, whenever we know something about the physics of the world it makes our representation learning task much simpler if we can run intervention experiments that test the causal dependencies between our latent variables (Squires et al., 2023; Ahuja et al., 2023).

We performed a small *proof-of-concept* experiment to test these predictions and report the results in Fig. 3. We train a standard VAE (with separate pose and, implicit, volume representation) on pseudo cryo-EM data and compare it to the same model but with the additional training step in Alg. 1. We refer to that model as `PoseVAE`. In Fig. 3B, we see that the additional penalty term in `PoseVAE` does, indeed, succeed at lowering the pose disentanglement metric (eq. 5). Inspecting the latent representations (Fig. 3E, middle), we observe that the pose is now fully confined to the pose variable. Moreover, we observe that the conformation space *z* is, itself, becoming more disentangled (Fig. 3E, right). Intuitively, this makes sense because less can go wrong now in encoding two instead of three variables into it. Quantitatively, this observation is confirmed by standard disentanglement metrics showing that `PoseVAE` achieves higher mean correlation coefficient (MCC) both across all latents (Fig. 3C, left) but also within the conformation latents *z* alone (Fig. 3, right). This is encouraging for the next task of disentangling the conformations.

## 4. DISENTANGLING CONFORMATIONS

Analogously to the previous section, in a second step, we propose a theoretical framework with metrics and benchmarks concerning the further disentanglement of the individual components inside the conformation vector *z* (Fig. 1). This addresses the essential challenge of interpretable cryo-EM conformational representations for heterogeneous reconstruction (Sec. 4.1). These benchmarks will help us measure true progress in the budding field of computational cryo-EM ([June 2023: Cryo-EM Heterogeneity Challenge](https://www.simonsfoundation.org/heterogeneity-in-cryo-electron-microscopy/)). Lastly, we discuss different methods to leverage recent development in nonlinear ICA that have the potential to build the next generation of cryo-EM models that get closer to the true answer (Sec. 4.2). These models may require future technological advancements such as temperature dependent cryo-EM (Bock and Grubmuller, 2022) or time-resolved single particle (X-ray) imaging (Shenoy et al., 2023a,b).

### 4.1 Evaluating Disentanglement of Independent Factors of Conformations

We consider the disentanglement of the individual components or dimensions inside the conformation space *z*. We propose a metric, in the form of a computational procedure, to evaluate whether the independent components of conformational variations are disentangled in the latent space. Essentially, this proposal relies on simulated data that exactly fulfills the generative model (eq. 1) which we assume for cryo-EM data. Having full control over the generative model is important, not only to measure progress, but also to simulate extended datasets (e.g., time-resolved imaging), because we know that only those future datasets with additional assumptions will, provably, allow progress in disentangling conformations. Otherwise, this challenge is hopeless (Locatello et al., 2019). Thus, in the ideal case, we have access to a good cryo-EM simulator *g*^*^, so we can just use this.

However, if we do not have a good cryo-EM simulator, we can just use the existing state-of-the-art model and see if we can recover its latents. More precisely, we can do the following: Train a regular cryo-EM model with the additional training loss (Alg. 1) that ensures that pose and conformation are disentangled. Then, check that the model is approximately, invertible. This is a common assumption in ICA theory (Hyvärinen et al., 2024) to make sure that the task of recovering the sources is well-defined. For this, we basically want to make sure that no two distinct points in conformation latent space *z*_1_ ≠ *z*_2_ would lead to the same volume representation *v*(*z*_1_) = *v*(*z*_2_). Once this is, approximately, validated we can use the model as a cryo-EM simulator. Intuitively, we now treat this first model as *ground truth* generator *g*^*^ := *g* and see if we can recover its latents *z*^*^ := *z*.

The procedure to evaluate and benchmark heterogeneous cryo-EM latent variable models would then be to assess how well they learn the same (up to equivalences) conformation latent space as the original model. Thus, we would effectively sample a ground truth pose *z*^*^ and some random pose *ϕ*^*^ and feed them into the *ground truth* model to obtain an image *x* = *g*^*^(*z*^*^, *ϕ*^*^) + *ϵ*. We then process this image with a candidate model *f*_*z*_(*x*) = *z* to obtain the learned conformation representation *z*. Consequently, we have to compare the two vector representations *z*^*^ and *z*. Depending on the equivalence class (∼_*C*_) that we are interested, there are many different metrics to assess how well *z*^*^ is disentangled in *z*. Intuitively, we want some kind of one-to-one correspondence between the two representations where changing a single entry in *z*^*^ corresponds to changing a single entry in *z*, and *vice versa*. Fortunately, this problem has been studied extensively in the machine learning subfield of disentangled representation learning (Bengio et al., 2013), with many proposed metrics and standardized benchmarks (Locatello et al., 2020a). We can build on those advances to get better quantitative measures on progress in heterogeneous cryo-EM reconstruction than volume based comparisons.

As an example metric measuring disentanglement, we will discuss the Mean Correlation Coefficient (MCC) (Hyvarinen and Morioka, 2017). Intuitively, we want each learned latent variable to be perfectly correlated (or anti-correlated, since sign flips do not compromise interpretability) with a single source variable. To measure this we can just compute the (absolute) correlation coefficient between all *ground truth* latents *z*^*^ and all learned latents *z*. To account for permutations, we have to solve a linear sum assignment with a permutation *σ* : {1, …, *K*} → {1, …, *K*}, which basically finds the best matching 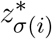 for each *z*_*i*_. The MCC is then, simply, the mean over those matches

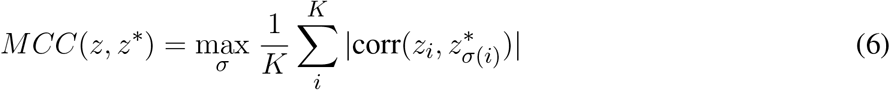

with corr() denoting correlation. Other metrics focus on decodability, or informational independence (Locatello et al., 2019) and there is no agreed-upon consensus on the optimal disentanglement metric. Thus, we simply report scores across all metrics – these can be further grouped by rank ordering to get overall model comparison scores (see Klindt et al., 2020).

### 4.2 Correcting Disentanglement of Independent Factors of Conformations

Let us now discuss the hardest task, i.e., finding the independent degrees of freedom that determine the conformation of a molecule (Fig. 1). This is a hard problem in the sense that, for a nonlinear function *g* (eq. 1), without any additional assumption it has been known for the last two decades that this is, practically, impossible (Hyvärinen and Pajunen, 1999). Moreover, the field of disentangled representation learning (Bengio et al., 2013) has spent multiple years proposing methods that were, ultimately, unidentifiable (Locatello et al., 2020a). Going forward, computational cryo-EM should learn from those lessons and avoid the same pitfalls. As a very basic example, if our conformation latent space has an isometric Gaussian prior, as in standard VAEs (Kingma and Welling, 2013), we can always perform a random rotation on the learned latents without changing the likelihood of the model (Hauberg, 2018). Thus, any direction in latent space may be representing the actual isolated change in conformation of the molecule. Fortunately, recent years have seen the development of different methods that solve the problem of nonlinear ICA (Hyvärinen et al., 2024). Below, we propose different approaches that are in technological reach (Sec. 4.2.1), or that make additional statistical assumptions that fit cryo-EM data (Sec. 4.2.2) or that integrate additional physical knowledge to constrain the problem (Sec. 4.2.3).

#### 4.2.1 Time-resolved Single Particle Imaging

If we had temporal data of conformation changes for the same molecule over time, we could start applying nonlinear ICA methods that depend on temporal autocorrelations of the sources (Hyvarinen and Morioka, 2017; Hälvä et al., 2021). Specifically, these methods operate under the assumption that we are able to record a time series like

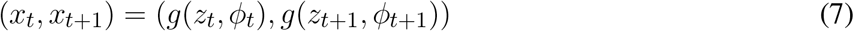

with temporal dependencies in the sense that

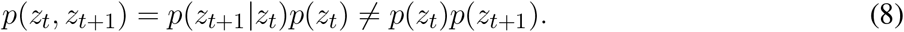

For instance, SlowVAE (Klindt et al., 2020) assumes that the transitions

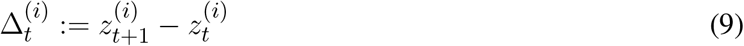

follow some sparse distribution, like a Laplace 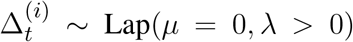, and the transitions are independent between components, i.e.,

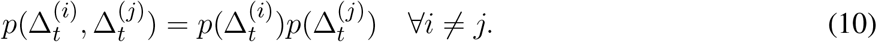

Klindt et al. (2020) showed that those assumptions are often verified on natural video data, which is important since making additional statistical assumptions to obtain identifiable models is only useful if those assumptions are, actually, aligned with the statistics of real world data.

Practically, such a model leads to a minimal modification to standard VAE training, where now the temporal difference of latents is also penalized to follow the specified transition distribution. However, the crucial difference in this learning paradigm is having access to temporal data 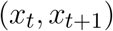. While this is not routinely feasible experimentally, efforts to develop time-resolved cryo-EM (Mäeots and Enchev, 2022; Lorenz, 2024) will eventually enable the direct observation of protein dynamics in the microseconds to seconds range, yielding datasets where each particle image will be associated to a timestamp that can be readily deployed in the modified modeling approach above.

We performed these disentanglement experiments in Fig. 4. In the first row, we have a demonstration of sparse transitions (drawn from a Laplace distribution) that show changes in some of the latent variables (*z*_0_, *z*_1_, *ϕ*). Below, we trained *N* = 50 models with and without temporal prior (Klindt et al., 2020) and measure seven typical disentanglement metrics (Locatello et al., 2019). We observe that in six out of seven of those metrics, the model with temporal prior, SlowVAE, does, indeed, achieve higher disentanglement scores. Further improvements could be achieved by including the pose loss from the previous section. However, this is already a promising proof-of-concept for future disentanglement of conformation latents based on temporal data.

**Figure 4.**
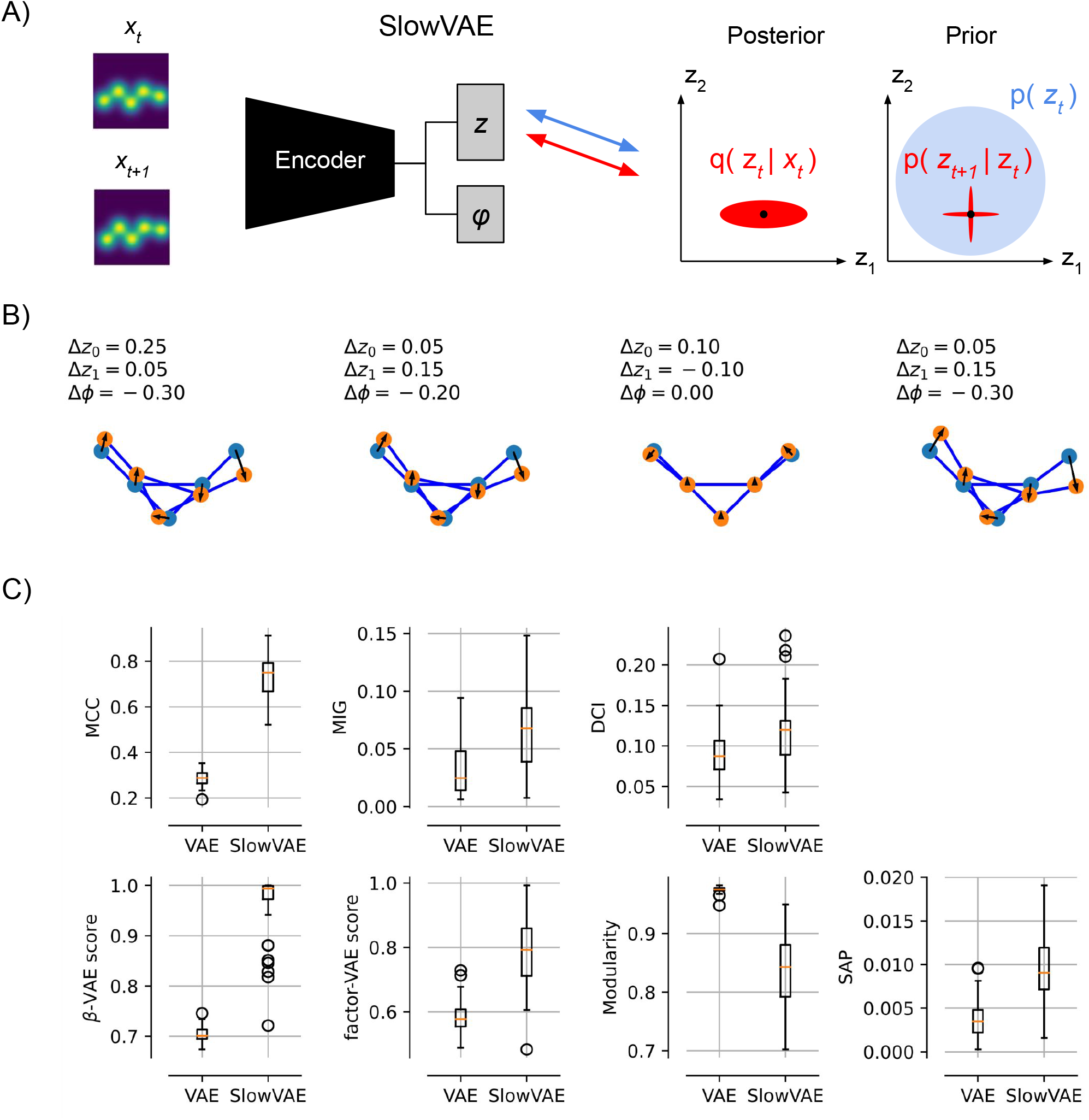
Temporal Conformation Disentanglement. **A)** Schematic of temporal data pairs (*x*_*t*_, *x*_*t*+1_) and minimal changes to VAE training. To obtain SlowVAE, all we need to do is change the conformation prior for the second time step to be a Laplace distribution centered around the posterior mean of the previous time step (Klindt et al., 2020). **B)** Example temporal transitions in the two conformation (*z*_0_, *z*_1_) and one pose *ϕ* latents, drawn from a Laplace distribution (default parameters from Klindt et al. (2020): data rate *λ* = 1, VAE (*β* = 1, *γ* = 10), VAE prior rate *λ*^*′*^ = 10. **C)** Most common disentanglement metrics, for details see (Locatello et al., 2019). In seven out of eight metrics we see that SlowVAE learns a more disentangled representation of the conformation latents than a regular VAE without any adaptations.

#### 4.2.2 Controlling the Boltzmann Distribution

The idea above applies to time-resolved experiments studying transient dynamics, triggered by some process such as mixing with a ligand or light excitation. Another class of experiments is concerned with steady-state dynamics where timestamps labels are not helpful. For those experiments, a potentially useful knob that could help solve the non-linear ICA problem is knowledge of the *temperature* associated with each particle in the dataset. The effect of cooling has been reviewed (Bock and Grubmuller, 2022) and different studies have used temperature to change the conformation distribution to obtain insights (Chen et al., 2019; Mehra et al., 2020). Experimentally, this could either be achieved by freezing grids with different cryogens, although a preferable approach would follow the development of thermochromic molecular probes able to report on the local temperature on the cryo-EM grid (Kortekaas and Browne, 2019). This way, precise temperature labeling of each particle in the dataset could be achieved.

Formally, manipulating the temperature *τ* would provide us with control over the Boltzmann distribution of molecular conformations

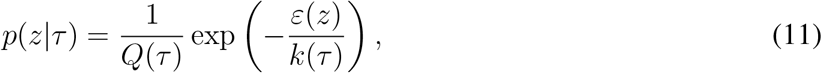

with *Q*(*τ* ) the canonical partition function and *ε*(*z*) the energy of being in conformation *z*. This conditional distribution, where we assume knowledge or experimental control over the temperature, maps onto the theoretical framework of iVAE (Khemakhem et al., 2020) with *u* = *τ* . Future theoretical investigations are needed to verify if the additional assumptions for their identifiability results are fulfilled in this setting.

However, to build intuition, we can walk through a thought experiment to see how control of the temperature can suffice to discover the independent degrees of freedom in molecular conformation changes (Fig. 5A). Assume, again a molecule with two degrees of freedom *z*_1_, *z*_2_ ∈ ℝ that both follow temperature-dependent normal distributions

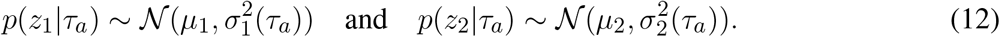

**Figure 5.**
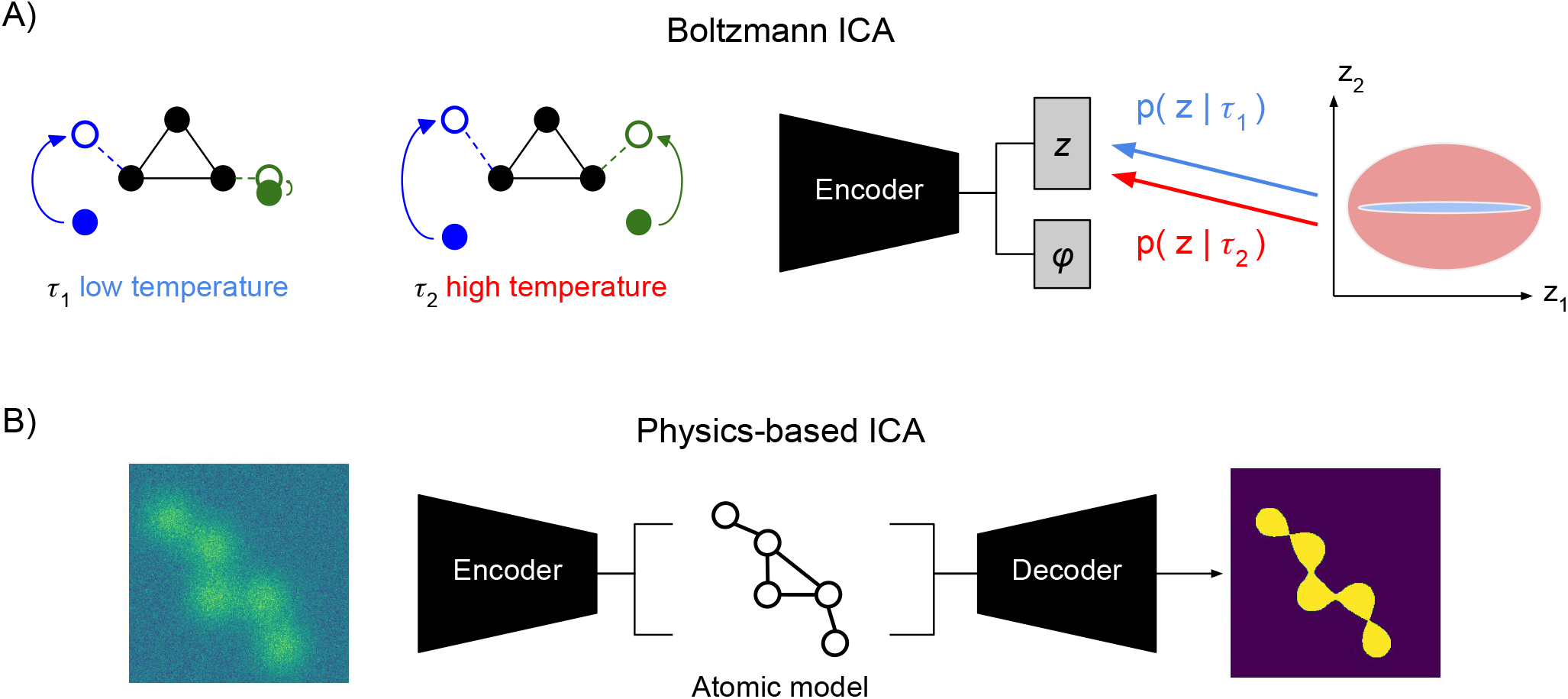
Alternative Conformation Disentanglement. **A)** Proof-of-concept illustration of **Boltzmann ICA** for the use of temperature as a conditioning variable (Khemakhem et al., 2020). At low temperature *τ*_1_ (left) only the left arm of the molecule shows significant movement. At high temperature *τ*_2_ both arms show significant movement. Conditioning the prior of the conformation latent with this information may allow identifying the distinct sources. **B)** Proof-of-concept illustration of **Physics-based ICA** using physics-based decoder with atomic models that assign mechanistic meaning to latent variables, e.g., in terms of atomic coordinates, (potentially, sparse) movement of volume (Punjani and Fleet, 2021), or local normal mode analysis (NMA) deformations (Nashed et al., 2022; Koo et al., 2023).

Now, suppose that at low temperature *τ*_*a*_, we only see variation in the first component *z*_1_ while the second component is nearly constant, i.e., 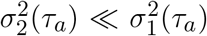. By contrast, at high temperature *τ*_*b*_ *> τ*_*a*_, we see that the second component also starts moving, i.e., 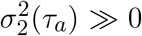. Thus, using temperature alone, we can successfully isolate the different degrees of freedom. Intuitively, this should make it possible to solve the disentanglement task. We could simply fit a model to the data at temperature *τ*_*b*_ with the additional constraint that the same model also has to be able to encode the data at temperature *τ*_*a*_, albeit, with only the first latent dimension *z*_1_. Whether those assumptions bare out in real molecules is not clear, yet, recording at different temperatures is within closer technological reach than time-resolved SPI.

#### 4.2.3 Atomic Models

While the previous two proposals require different data, we may also hope to make progress with different models. We saw how the implicit volume representation needs additional care to disentangle pose and conformation information (section 3.1). Imbuing the generative model with physics inspired structure, allowed us to separate pose from conformation (section 3.2). Maybe, even more physics can help us solve the harder problem of finding the conformational degrees of freedom. In particular, if we replace the highly expressive implicit volume representation *v* with an atomic model (Zhong et al., 2021b; Rosenbaum et al., 2021; Nashed et al., 2022; Koo et al., 2023), then the pose latent variable *z* will have to encode how an atomic reference structure (maybe the mode of the conformation space) is deformed into an observed conformation (Fig. 5B).

One existing work by Punjani and Fleet (2021) proposes to learn a convection field that deforms a reference volume. This comes with the elegant property of *volume preservation* which is not always the case in implicit conformation representations, but obeys our knowledge of the underlying physics. However, the learned convection fields as well as the reference volume model are still over-parameterized compared to an ideal atomistic model with movement vectors for each atom. The problem in building smaller and more constraint models is that modern machine learning methods have, to some extent, proven so powerful because they allow heavily overparameterized hypothesis classes that still generalize well beyond the training data. The question then becomes whether we can combine the best of both worlds, i.e., the non-convex optimization and generalization properties of deep neural networks (e.g., for implicit volume or convection representations) with the physical detail of constraint atomistic models (Zhong et al., 2021b; Rosenbaum et al., 2021; Nashed et al., 2022; Koo et al., 2023).

## 5 DISCUSSION

In recent years, the integration of computational models, particularly VAEs (Kingma and Welling, 2013), has revolutionized research across various natural sciences, including cryo-EM (Zhong et al., 2021a). This perspective piece underscores the critical importance of understanding conformational latent spaces in cryo-EM by drawing on cutting-edge theoretical advancements in identifiable nonlinear ICA (Hyvärinen et al., 2024). By bridging the gap between theoretical frameworks and practical applications in cryo-EM, we are suggesting a significant advance in the interpretability and utility of latent variable models. Furthermore, our study advocates for the adoption of better quantitative measures to assess progress in heterogeneous cryo-EM reconstruction, transcending traditional volume-based comparisons. This aligns with recent initiatives such as the [Cryo-EM Heterogeneity Challenge](https://www.simonsfoundation.org/heterogeneity-in-cryo-electron-microscopy/) emphasizing the need for refined evaluation metrics to accurately gauge advancements in this field. Nevertheless, this work is merely an opinion piece and *proof-of-concept* demonstration. Significant technical (e.g., time resolved SPI) and engineering challenges (e.g., identifiable nonlinear ICA models that work in low SNR regimes) lie ahead on this path towards interpretable cryo-EM conformations spaces.

The history of ICA’s emergence in the 1990s, and in particular its early adoption in neuroimaging (McKeown et al., 1998), shows its capacity to evolve into a cornerstone of data-based modeling. This trend, moving even further away from hypothesis-driven research (Friston, 1998) toward data-centric approaches (Beckmann et al., 2005), also underscores the importance of incorporating principles like nonlinear ICA to ensure meaningful model outputs. While modern machine learning techniques, including VAEs or nonlinear dimensionality reduction methods such as t-SNE (Van der Maaten and Hinton, 2008) or UMAP (McInnes et al., 2018), have become ubiquitous in data-based modeling, they often overlook source recovery, i.e., identifiability considerations. To fully harness the potential of latent spaces, it is paramount to ensure their alignment with meaningful representations of the underlying data.

In conclusion, our approach integrates nonlinear ICA principles into the development and analysis of cryo-EM latent variable models, ensuring more interpretable representations that encapsulate the intrinsic structure of the data. Unlocking latent spaces aligned with the underlying fundamental factors governing complex phenomena is pivotal for gaining deep insights into biological processes, expediting drug discovery, and facilitating targeted interventions. This progress extends beyond cryo-EM, resonating with diverse scientific disciplines such as computer vision, natural language processing, and generative modeling, where (VAE) latent spaces play a pivotal role in data representation and the generation of new scientific hypothesis as part of initiatives such as AI4Science. Our interdisciplinary approach, embracing nonlinear ICA and disentanglement models, holds promise in generating meaningful representations that *carve nature at the joints*, thereby propelling transformative discoveries.

## AUTHOR CONTRIBUTIONS

D.K. N.M. and F.P. conceived the study. N.M. and F.P. provided guidance, feedback and the intellectual environment of this work. A.H. N.M. and D.K. formalized the problem and provided the link between cryo-EM and ICA. D.K. and A.L. developed the pose disentanglement metrics. D.K. performed all experiments and made the figures. All authors helped writing the manuscript.

## ACKNOWLEDGMENTS

This work was inspired by discussions at the Flatiron CryoEM Summer Workshop 2023. Specifically, we acknowledge Yifan Cheng’s determination to really understand conformational latent spaces. We hope that the present work will help the field of computational cryo-EM to this next level where they can provide truly novel insights into Biology.

## APPENDIX

### 5.1 Pose and Conformation Disentanglement Metrics

We consider disentanglement evaluation metrics for all three learned functions, i.e., the volume *v*(*z*), the conformation encoder *f*_*z*_(*x*) and the pose encoder *f*_*ϕ*_. For each function, we will consider two metrics that measure their consistency (i.e., does it model the latent it is supposed to model?) and their invariance (i.e., is it invariant to changes in other latents?). Starting with the *volume metrics*, we measure

*1. Volume consistency*, i.e., how accurately the learned volume *v*(*z*) changes the conformation:

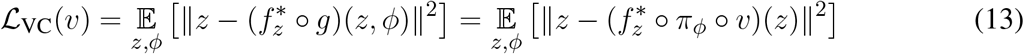

*2. Volume pose invariance*, i.e., how accurately the learned volume *v*(*z*) changes *only* the conformation:

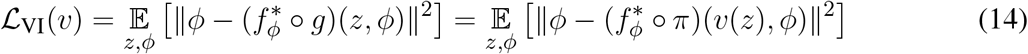

where, ideally, one would use the oracle encoder *f*^*^(*x*) = argmax_*z,ϕ*_ *p*(*x*|*z, ϕ*) *p*(*z, ϕ*). In practice, we can just use the current encoder *f* (*x*) = (*f*_*z*_(*x*)||*f*_*ϕ*_(*x*)) and optimize it as well. Again this can be split into two metrics (consistency and invariance), both for the *conformation encoder f*_*z*_

*1. Conformation-encoder consistency*, i.e., how accurately the conformation encoder *f*_*z*_ recovers any conformation, independent of pose:

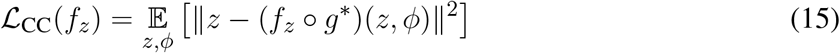

*2. Conformation-encoder pose invariance*, i.e., how invariant the conformation encoder *f*_*z*_ is to pose perturbations:

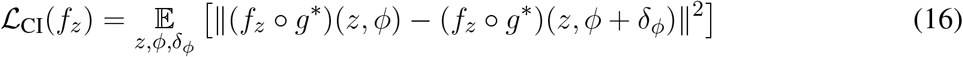

as well as for the *pose encoder f*_*ϕ*_

*1. Pose-encoder consistency*, i.e., how accurately the pose encoder *f*_*ϕ*_ recovers any pose, independent of conformation:

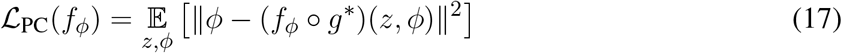

*2. Pose-encoder conformation invariance*, i.e., how invariant the pose encoder *f*_*ϕ*_ is to conformation perturbations:

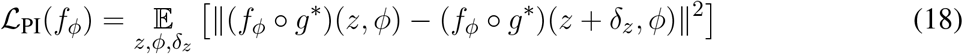

where, again, ideally we would like to use the *ground truth* generator *g*^*^, but we can also just use the current learned decoder. All of these six metrics can be evaluated, in a supervised way, over a sufficient number of randomly sampled conformations *z*, poses *ϕ* and perturbations (*δ*_*z*_, *δ*_*ϕ*_).

## Footnotes

1 Precisely, writing *f* (*x*) = *Wx*, ∀*z* ∈ *Ƶ* : (*f* ◦ *g*)(*z*^*^) ∼_*C*_ *z*^*^ ⇔ *WA* = *DP* where *D* is a diagonal matrix and *P* is a permutation.

2 In the common framework of VAEs, *f*_*z*_ (*x*) = *µ*_*z*_ (*x*) could be defined as the mean of the variational posterior; in an auto-decoding framework this could be the MAP outcome of inference, i.e., *f*_*z*_ (*x*) = argmax_*z*_ *p*(*x*|*z*)*p*(*z*) (see appendix XXX).

